# Antagonistic roles of NOT1 paralogues in the timing of gene expression in *Plasmodium falciparum*

**DOI:** 10.1101/2020.02.12.945477

**Authors:** Ying Liu, Ragini Rai, Lei Zhu, Changqing Zhang, Frances Rocamora, Mark Featherstone, Zbynek Bozdech

## Abstract

NOT1 is the scaffold of the CCR4-NOT complex, a highly conserved multi-protein complex that regulates gene expression in eukaryotes. As opposed to most eukaryotes in which NO1 is encoded by a single gene, malaria parasites, *Plasmodium falciparum,* carry two NOT1 paralogues, PfNOT1.1 and PfNOT1.2. Here we showed that the two PfNOT1 proteins function as mutually exclusive scaffolds within the PfCCR4-NOT protein complexes that are abundantly located in the parasite cytoplasm. Intriguingly, the two PfNOT1 paralogues appear to have directly opposing functions in regulation of mRNA abundance across the *P. falciparum* IDC, in which PfNTO1.1 and PfNOT1.2 induces and suppresses transcript abundance during their active transcription, respectively. Targeted disruption of either of the *PfNOT1* gene causes defective growth and lower invasion rates presumably due to the deregulation the *P. falciparum* IDC transcriptional cascade. We also demonstrate that the regulatory function of both PfNOT1.1 and PfNOT1.2 are related to another PfCCR4-NOT subunit, *PfCaf1,* which indicates their activity during post-transcriptional regulation. Indeed RNA decay studies suggest the active role of both PfNOT1 proteins in regulation of mRNA stability in a directly opposing manner.

**Author summary:** CCR4-NOT complex is a highly conserved multi-protein complex that regulates gene expression in eukaryotes. NOT1 serves as the scaffold of the complex and plays important roles in gene regulation both transcriptionally and post-transcriptionally. As opposed to other eukaryotes, *P. falciparum* encodes two paralogues of PfNOT1, raising the question as to the significance to possess an additional copy of PfNOT1 in the parasite. Here we described antagonistic regulatory functions of two PfNOT1 paralogues in gene expression during the 48-hour intraerythrocytic developmental cycle. We also reported that their regulatory functions are predominantly post-transcriptional and proposed a model in which distinct PfCCR4-NOT complexes defined by mutually exclusive PfNOT1 scaffolds differentially regulate PfCAF1 function in mRNA decay. This study highlights the importance of post-transcriptional regulation in *P. falciparum* and provides novel insights into mechanisms of gene regulation in this organism. The unique presence of two PfNOT1 paralogues may also open avenues for the development of new drug targets for anti-malarial control.

## Introduction

*Plasmodium falciparum* is an *Apicomplexan* parasite responsible for the most lethal form of malaria and is therefore a major global health concern [1]. The lifecycle of this protozoan parasite consists of a sexual stage in the *Anopheles spp.* mosquito host and an asexual stage in the human host [2]. In a human red blood cell, the parasite passes through three morphological stages – ring, trophozoite and schizont – across the 48-hour intraerythrocytic developmental cycle (IDC), eventually releasing up to 32 merozoites ready to invade red blood cells in a subsequent round of infection [3]. During one round of the IDC, the expression of more than 50 % of genes displays a continuous cascade with transcript levels peaking at the point when their protein products are needed, leading to the “just in time” model of gene expression [4]. Hence, gene expression is highly regulated across the IDC at multiple levels, ranging from epigenetic regulation, transcription and post-transcriptional regulation.

While the basic mechanism of eukaryotic transcription and general transcription components are conserved in *P. falciparum* [5, 6], most TBP-associated factors (TAF) are absent [6] and only a third of the expected number of canonical eukaryotic transcription activating proteins (TAP) were predicted in the parasite genome [7]. The presence of 27 *Apicomplexan*-specific transcription factors, ApiAP2, may compensate for the reduction in the number of conventional transcription components [8, 9]. These 27 ApiAP2 proteins display distinct expression profiles covering the entire IDC [8] and their recognition motifs are located in the promoter regions of most genes whose expression profiles are positively correlated to expression of the corresponding ApiAP2 gene [9]. In addition to transcriptional initiation, there is mounting evidence that multiple processes of post-transcriptional regulation are also critical for regulation of mRNA abundance through the *P. falciparum* IDC [10–12]. This is supported by the abundant presence RNA-binding proteins [13, 14] and RNA degradation components [15] in the *P. falciparum* genome. The includePfALBA1 [16], PfCAF1 [17] in *P. falciparum* and PyCCR4 [18], PyALBA4 [19] in rodent-specific *P.yoelii* all of which functions were directly implicated in regulation of transcript levels. Moreover, variability of mRNA half-lives across the IDC observed by several experimental approaches also suggested a significant contribution of mRNA decay in regulation of gene expression in malaria parasites [20]. Taken together, both transcriptional regulation and post-transcriptional regulation play critical roles in modulating transcript abundance in *P. falciparum*.

CCR4-NOT is a highly conserved proteins complex involved in multiple steps of gene expression regulation across the entire eukaryotic kingdom [21], present in both the nucleus and cytoplasm [22, 23]. Core subunits include NOT1, NOT2, NOT3/5 paralogues, NOT4, CAF1, CCR4, CAF40, yeast-specific CAF130 and *Drosophila*/mammalian-specific NOT10 and NOT11 [21]. Structurally, NOT1 is the largest subunit of the CCR4-NOT complex and acts as a scaffold to recruit other subunits [24]. The N-terminal domain of NOT1 interacts with NOT11 and NOT10 in *Drosophila* and mammals [25]. The central MIF4G domain associates with CAF1, which bridges CCR4 to the whole complex [24]. The association of CAF1 and CCR4 to the complex is required for their functions as deadenylases to shorten the mRNA poly(A) tail in yeast and *Drosophila* [26–28], the key step in mRNA degradation[29]. The DUF3819 domain of NOT1 interacts with CAF40 [30]. The C-terminal NOT1 domain interacts with NOT2 and NOT3/5 paralogues [31], which associate with transcription factors [32, 33] and histone modifications [34, 35]. The NOT1 domain also interacts with NOT4 [31], an E3 ubiquitin ligase involved in protein degradation [36, 37] and required for proteasome integrity in yeast [38].

In spite of its broad involvement in multiple levels of eukaryotic gene expression, only limited information exist about the biological function of the CCR4-NOT complex in malaria parasites; particularly its significance for the progression of the parasite life cycle. Initially it was shown that targeted gene disruption of *PfCaf1* affected expression of 1,031 genes, including many invasion-related genes, which leads to the release of premature merozoites in *P. falciparum* [17]. In *P. yoelii*, PyCCR4 appears to regulate gene expression in gametocytes while disruption of *PyCaf1* or *PyCcr4* results in a reduction of male gametocyte activation [18]. Moreover, PyCCR4 was show to associate with some transcripts that are dysregulated in the parasite lacking *PfCcr4*A potentially via its intrinsic DNA nuclease activity that is dependent on Cu^2+^ [39]. These studies demonstrate the importance of the presumptive CCR4-NOT complex in *Plasmodium* parasite at multiple stages. The NOT1 subunit has been identified as a negative transcription regulator in yeast [22], although a positive effect on transcription has also been reported [40]. The association of yeast NOT1 with several transcriptional components, including TBP [41], TAFs [42] and the RNAPII elongation complex [43, 44] reinforces a role for NOT1 in transcription. Post-transcriptionally, NOT1 interacts with multiple RNA-binding proteins, including tristetraprolin [45], eIF4E-binding protein 4E-T [46], RNA helicase Xp54 [47] and m6A reader protein YTHDF [48], involved in mRNA degradation, mRNA decapping and translation repression in multiple organisms. Moreover, NOT1 in mouse germ cells interacts with the RNA-binding protein Nanos2, which is speculated to promote the localization of CCR4-NOT complex to P-bodies, the center of RNA degradation [49]. The structural role as the scaffold of the CCR4-NOT complex and implication in both transcriptional and post-transcriptional regulation makes NOT1 an ideal focus to investigate the presumptive PfCCR4-NOT complex and gene regulation in *P. falciparum*.

Identically to all characterized *Plasmodium* species*, P. falciparum* parasites encode two paralogues of PfNOT1 located on chromosome 11 (termed PfNOT1.1) and chromosome 14 (termed PfNOT1.2), raising the question as to the composition of tentative CCR4-NOT complex and its regulatory role in gene expression. In this study, we demonstrate that while the overall structure of the *P. falciparum* CCR4-NOT complex is highly conserved, PfNOT1.1 and PfNOT1.2 function as mutually exclusive scaffolds. Intriguingly, PfNOT1.1 and PfNOT1.2 appear to have antagonistic functions in regulation of gene during the IDC progression, presumably via differential interaction of the CCR4-NOT complex with other regulatory factors. Our results suggest that the regulatory functions exerted by the two PfNOT1 paralogues is mediated by factors of mRNA decay.

## Results

### 1. Four conserved CCR4-NOT subunits are localized in the cytoplasm

Most core subunits of the CCR4-NOT complex are conserved in the *P. falciparum* genome, including PfNOT2, PfNOT4, PfNOT5, PfCAF1 and PfCAF40 [50] (**See Table. S1**). However, unlike other eukaryotes studied to date, the malaria parasites have two PfNOT1 and four PfCCR4 paralogues and conversely do not harbor homologues of CAF130, NOT10 and NOT11. While the loss of CAF130, NOT10 and NOT11 could be attributed to the overall genome streamlining documented in *Plasmodium* species [50], the duplications of the PfNOT1- and the PfCCR4-coding genes are unique amongst eukaryotic organism. Canonically, NOT1 functions as a structural scaffold protein in the CCR4-NOT complex and as such is encoded by a single gene copy in the vast majority of eukaryotic species [21]. Nonetheless, for both PfNOT1.1 and PfNOT1.2, there is a high degree of evolutionary conservation with the presence of the MIF4G, DUF3819 and NOT1 domains (**Fig. S1A and S1B)**. There are apparent insertions of unique amino acid stretches within the MIF4G domain and the NOT1 domain in PfNOT1.2, exclusively. Sequence insertions in highly conserved domains have been reported in *Plasmodium* proteins, suggesting their rapid evolution and diversification [51]. This suggests that PfNOT1.1 may reflect the ancestral gene copy, while PfNOT1.2 is a product of recent duplication and thus may have diverged from its original function presumably as an alternative scaffold in CCR4-NOT complex.

**Table S1.** Predicted CCR4-NOT subunits in *P. falciparum*.

| *P. falciparum* | Gene ID | Yeast | Human | *Drosophila* |
| --- | --- | --- | --- | --- |
| PfNOT1.1<br>PfNOT1.2 | PF3D7_1103800<br>PF3D7_1417200 | CDC39 | CNOT1 | Not1 |
| PfNOT2 | PF3D7_1128600 | CDC36 | CNOT2 | Regena (Rga) |
| PfNOT4 | PF3D7_1235300 | MOT42/SIG1 | CNOT4 | Not4 |
| PfNOT5 | PF3D7_1006100 | NOT3<br>NOT5 | CNOT3 | Not3 |
| PfCAF1 | PF3D7_0811300 | POP2 | CNOT7<br>CNOT8 | Pop2 |
| PfCCR4.1<br>PfCCR4.2<br>PfCCR4.3<br>PfCCR4.4 | PF3D7_0519500<br>PF3D7_1363500<br>PF3D7_0107200<br>PF3D7_0319200 | CCR4 | CNOT6L<br>CNOT6 | Twin |
| PfCAF40 | PF3D7_0507600 | CAF40 | CNOT9 | Rcd1 |
|  |  | CAF130 |  |  |
|  |  |  | CNOT10 | Not10 |
|  |  |  | CNOT11 | Not11 |

The CCR4-NOT core subunits identified in yeast, human and *Drosophila* are listed. Their counterparts in *P. falciparum* were predicted by searching the PlasmoDB database and blasting protein sequences of each subunit in the *P. falciparum* proteome.

To explore this further, we created independent transgenic parasites in which the endogenous allele of *PfNot1.1* and *PfNot1.2*, as well as *PfCaf1* and *PfNot4* were joined with HA epitopes by single crossover recombination (**Fig. S2A**). The correct insertion of the HA epitopes was verified by genotyping PCR for all four transgenic lines, including PfNOT1.1::HA, PfNOT1.2::HA, PfCAF1::HA and PfNOT4::HA (**Fig. S2B-E**). Western blotting analysis indicated that the bulk of PfNOT1.1 and PfNOT1.2 is present predominantly in the cytoplasm, while PfCAF1 and PfNOT4 could be found in both the cytoplasm and nucleus (**Fig. 1A-D**). This was supported by the subsequent immunofluorescent assay showing the four proteins present in the cytoplasm as a punctuate pattern (**Fig. 1E-H**). These cytosolic puncta did not overlap with aldolase or DAPI, whcih is consistent with previous reports for the P. yoelii orthologue, PyCCR4 [18] and the other three PfCCR4 paralogues [39]. This intracellular localization is consistent with other eukaryotic species suggesting the function of PfCCR4-NOT complex in posttranscriptional regulation via interactions with mRNA in the parasite cytoplasm.

**Figure 1.**
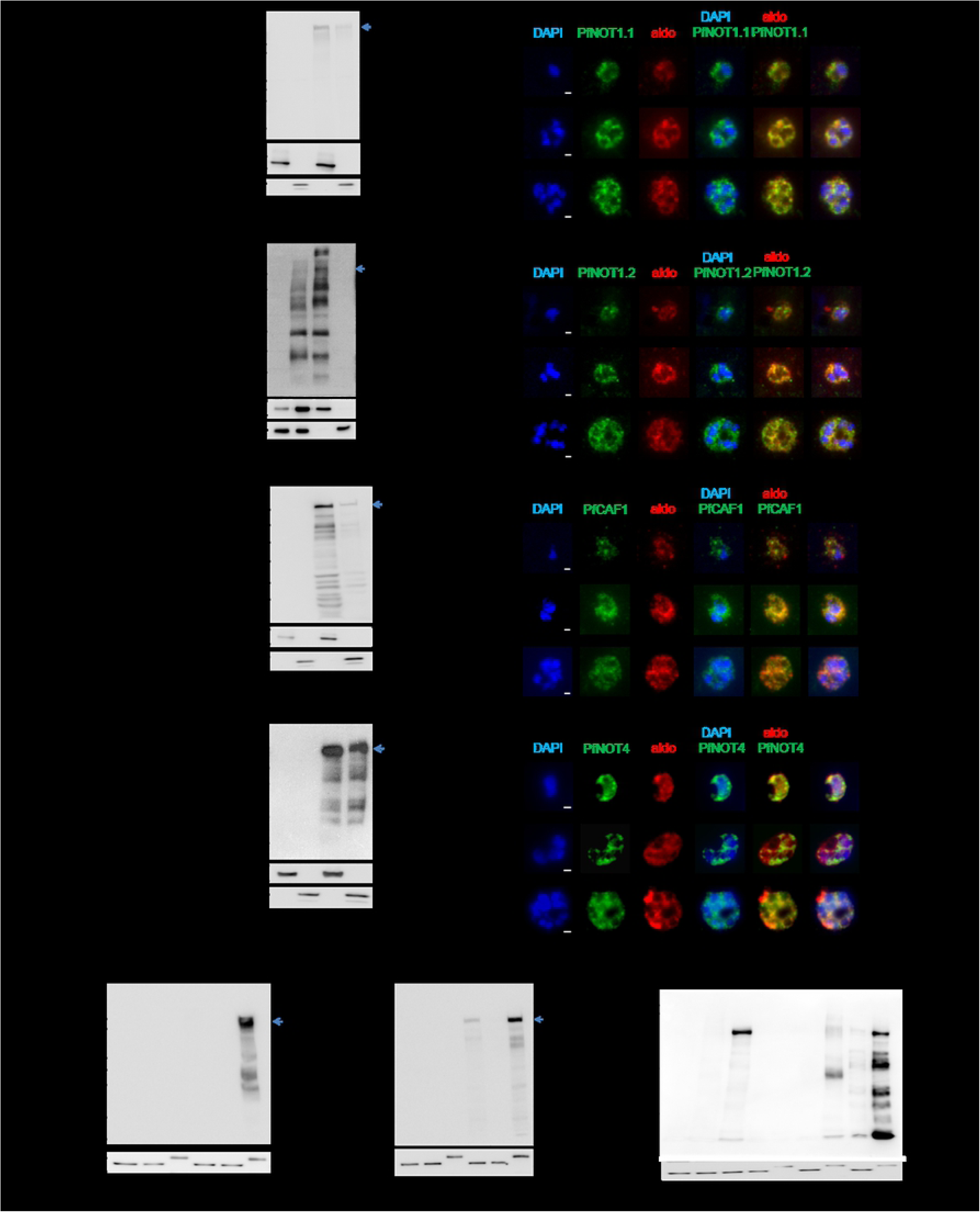
Four conserved CCR4-NOT subunits are localized in the cytoplasm. Expression of HA-tagged PfNOT1.1 **(A)**, PfNOT1.2 **(B)**, PfCAF1 **(C)** and PfNOT4 **(D)** was tested by western blotting using anti-HA antibody of total lysates, cytoplasmic lysates (Cyto) scored by aldolase and nuclear lysates (Nuc) scored by histone H3. The location of the full-length protein is indicated by arrows. Four proteins were detected predominantly in the cytoplasmic lysates. (**E-H**) Four studied proteins were visualized as cytosolic puncta by the immunofluorescent assay using anti-HA antibody at the ring, trophozoite and schizont stages. Nuclei were stained with DAPI and aldolase immunoreactivity was used as a cytoplasmic maker. Scale bars are 1 micron. Immunofluorescent assay on the parental strain served as a negative control (Fig. S2F in supplemental figures). **(I-K)** Immunoprecipitation of HA-tagged PfNOT1.1, PfNOT1.2, PfCAF1 and PfNOT4 was verified by western blotting. HA-tagged proteins were detected in the input and eluent but not the unbound supernatant of each HA-tagged transgenic line by anti-HA antibody. Immunoprecipitation of the parental strain lysate (par) served as a negative control. Comparable signals of actin verified similar loading of each parasite sample. The sizes of the ladder are given at the left.

### 2. PfNOT1.1 and PfNOT1.2 function as mutually exclusive scaffolds of the P. falciparum PfCCR4-NOT complexes

To further investigate the composition of the PfCCR4-NOT complex, we carried out co-immunoprecipitations coupled with mass spectrometry on the four HA-tagged proteins. Prior to the mass spectrometry, we confirmed the efficiency of immunoprecipitation of HA-tagged proteins using western blotting, showing their strong abundance in the eluent fractions (**Fig. 1I-K).** Mass spectrometry analyses of the eluent fractions detected 30 proteins for PfNOT1.1, 20 proteins for PfNOT1.2, 244 proteins for PfCAF1 and 45 proteins for PfNOT4 in two biological replicates (see full protein lists in **Table S2).** The co-immunoprecipitation studies confirmed physical interaction of PfCAF40 with PfNOT1.1, PfNOT1.2, PfCAF1 and PfNOT4; PfCAF1 with PfCCR4; and either of the PfNOT1.1 or PfNOT1.2 with PfCAF40 and PfCAF1, respectively. On the other hand PfNOT4 failed to co-immunoprecipitate PfNOT1.1 and PfNOT1.2 **(Table 1, top panel)**, which is consistent with the transient character of this subunit with the CCR4-NOT complex previously demonstrated in *Drosophila* [52], human [53] and *P.yoelii* [18]. However, our data showed that PfNOT4 does associate with the PfCCR4-NOT complex given its interaction with CAF40 and possibly CAF1. Overall, this analysis supports the presence of the four studied proteins in the PfCCR4-NOT complex, demonstrated by their interactions with each other and other canonical subunits.

**Table 1.**
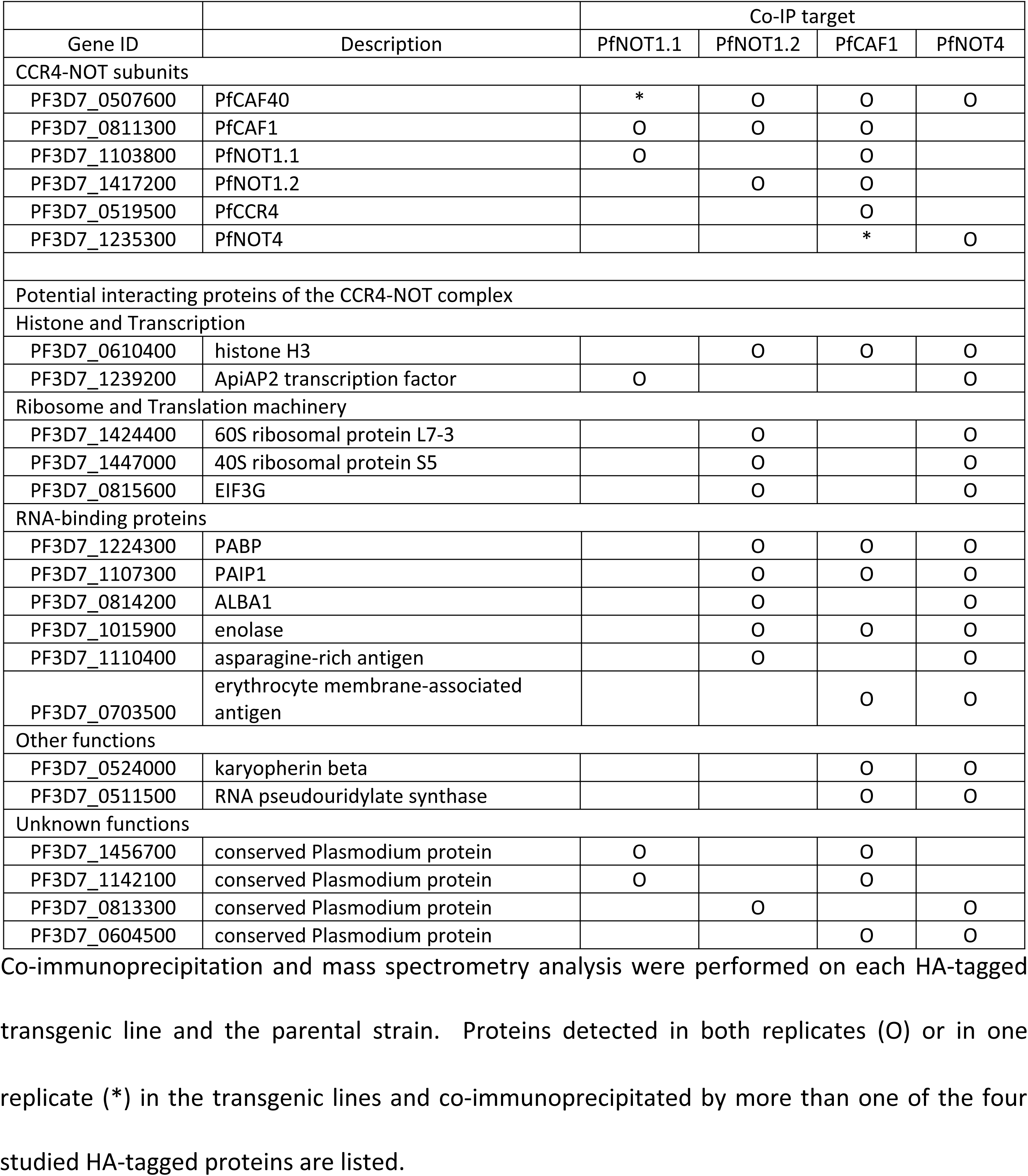
Potential interacting proteins of PfNOT1.1, PfNOT1.2, PfCAF1 and PfNOT4.

Interestingly the two PfNOT1 paralogues did not co-immunoprecipitate each other. This is consistent with the model that both PfNOT1.1 and PfNOT1.2 function as scaffolds in the PfCCR4-NOT complexes in the mutually exclusive manner; given that the structure of the eukaryotic CCR4-NOT complexes contains a single NOT1 subunit. This is also consistent with the exclusivity of the other (than CCR4-NOT subunit) proteins that were co-immunoprecipitated by each of the PfNOT1 paralogues (**Table 1, bottom panel**). Particularly, PfNOT1.1 co-immunoprecipitated proteins involved in transcription (chromodomain helicase DNA-binding protein 1 homolog PF3D7_1023900, acetyltransferase PF3D7_1323300 and ApiAP2 transcription factor PF3D7_1239200), Maurer’s clefts components (protein phospharase PP1 PF3D7_1414400, exported protein PF3D7_0501000 and pyruvate kinase PF3D7_0626800) and RNA-binding (polyadenylate-binding protein PF3D7_1360900). PfNOT1.2 co-immunoprecipitated proteins involved in RNA-binding (polyadenylate-binding protein PABP PF3D7_1224300, PAIP1 PF3D7_1107300 and ALBA1 PF3D7_0814200), translation (4 ribosomal proteins and EIF3G PF3D7_0815600) and histone composition (histone H2B and histone H3) (see full protein lists in **Table S2**). Overall, the associations of proteins involved in transcription, translation and RNA-binding suggests the role of the PfCCR4-NOT complexes in regulation of *P. falciparum* gene expression that is orthologous to other eukaryotes [29]. However, the differential protein associations between the PfNOT1.1 and PfNOT1.2-containig complexes support the notion that the two PfNOT1 paralogues modify the overall activity of the PfCCR4-NOT complex presumably giving it distinct sets of functions in gene regulatory processes.

### 3. Targeted disruption of PfNot1.1 or PfNot1.2 impairs parasite proliferation and invasion

To explore this, we carried out targeted gene disruptions for both *PfNot1.1* and *PfNot1.2* in independent parasite lines. First, we disrupted the *PfNot1.1* coding region by inserting a human dihydrofolate reductase (*hDHFR*) cassette via double crossover recombination (ΔPfNot1.1) (**Fig. 2A**). The cloned parasite line was obtained by limiting dilution and the construct insertions was verified by PCR-based genotyping (**Fig. S3A**). Finally, Northern blot analysis showed a complete absence of PfNOT1.1 transcript (**Fig. 2B**). A similar strategy of gene disruption was, however, unsuccessful for *PfNot1.2* after three independent transfection experiments (**Fig. S3B**), suggesting that PfNOT1.2 might be essential for parasite viability. Thereafter, we created a conditional protein knockdown parasite line to express PfNOT1.2 fused with an *E.coli* DHFR destabilizing domain (ecDDD) at the C-terminus (ddΔPfNot1.2) (**Fig. 2C**). In this system, the transgenic protein is stabilized by the inclusion of the trimethoprim (TMP) ligand in the culture medium. Indeed, the ecDDD insertion was achieved by single crossover recombination and confirmed by genotyping PCR (**Fig. S3C**). Using this strategy, we achieved a significant reduction of PfNOT1.2 expression at the schizont stage after removing TMP for three consecutive IDCs (**Fig. 2D** and **Fig. S3D).**

**Figure 2.**
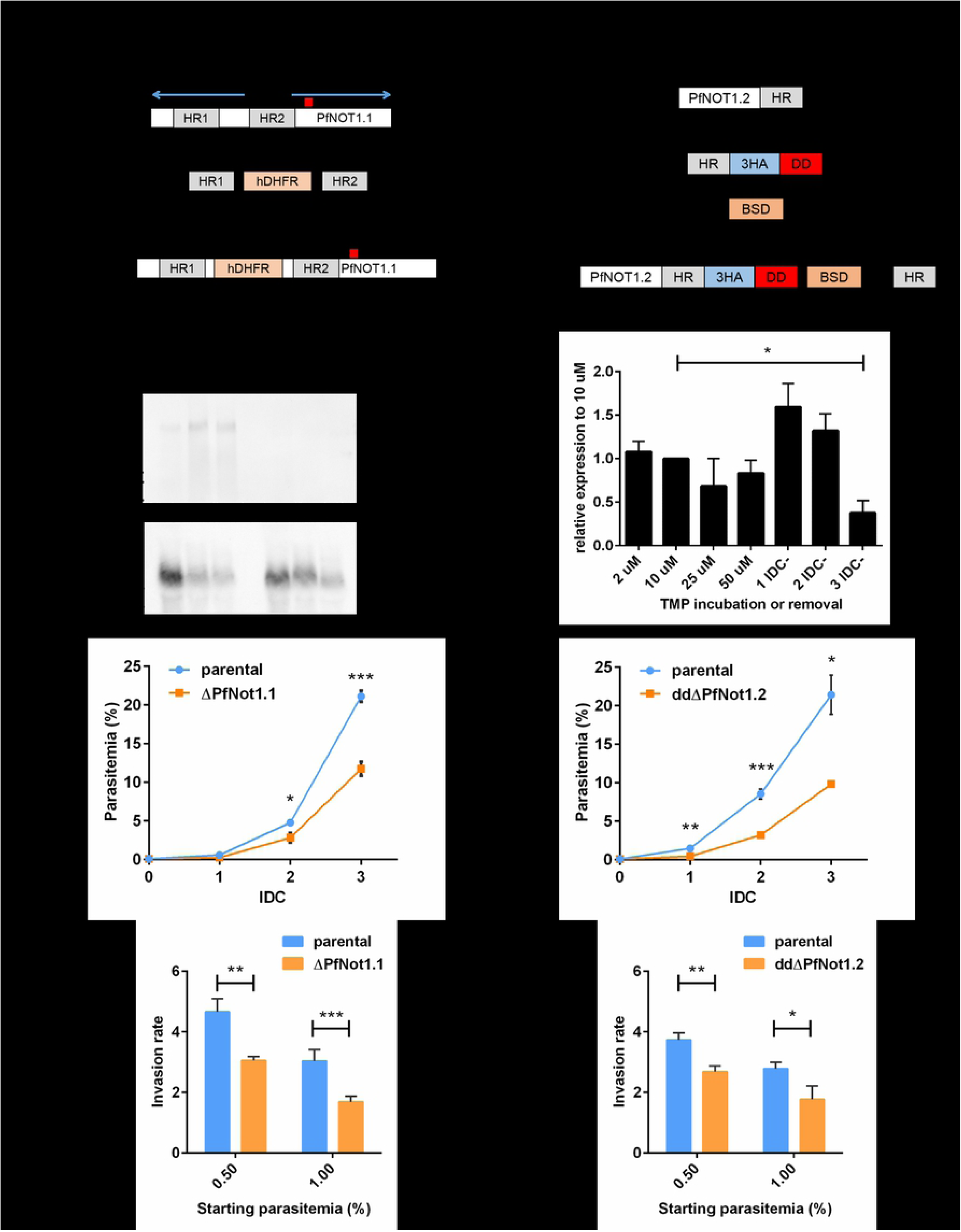
Targeted disruption of *PfNot1.1* or *PfNot1.2* impairs parasite proliferation and invasion. **(A)** The endogenous *PfNot1.1* was disrupted by the *hDHFR* cassette by double crossover recombination. **(B)** Northern hybridization analysis detected *PfNot1.1* transcript in the parental strain but not ΔPfNot1.1 at the ring (R), trophozoite (T) and schizont (S) stages with the probe shown as red boxes in Fig. 1A. The loading control is Serine-tRNA ligase transcript. **(C)** HA epitopes and an ecDDD destabilizing domain were fused to PfNOT1.2 in the C-terminus by single crossover recombination. **(D)** Expression of tagged PfNOT1.2 was detected by western blotting in biological triplicates. Intensities were quantified by ImageJ and normalized to actin expression level and then to the expression level in 10 μM TMP, the concentration employed to maintain the culture. Comparable protein intensities for different concentrations of TMP ligand indicates that 2 μM TMP is sufficient to stabilize the ecDDD. Tagged PfNOT1.2 was conditionally knocked down significantly after removing TMP for three IDCs (p < 0.05). **(E, F)** Parasitemia was counted in synchronized cultures with biological triplicates across three IDCs. Parasite proliferation rate of ΔPfNot1.1 and ddΔPfNot1.2 was significantly lower compared to that of the parental strain. **(G, H)** Two transgenic lines showed significantly lower invasion rate compared to that in the parental strain in biological triplicates. All data are represented as mean ± standard deviation from three biological triplicates and statistic test was performed with p < 0.05 (*), p < 0.01 (**) and p < 0.001 (***).

Disruption of *PfNot1.1* and conditional knockdown of *PfNot1.2* both resulted in significantly lower proliferation rates across three consecutive IDCs (**Fig. 2E, 2F**). In particular, ΔPfNot1.1 and ddΔPfNot1.2 exhibited considerable decreases in invasion rate with more moderate defects in ddΔPfNot1.2 (**Fig. 2G, 2H**), possibly due to incomplete knockdown. This proliferation defect is not associated with the numbers of merozoites produced per schizont by each transgenic line that was comparable to their parental line (**Fig. S3E and S3F**). Moreover, maximum likelihood-based statistical analysis of the IDC transcriptomes showed no significant differences in the progression of ΔPfNot1.1 and ddΔPfNot1.2 (**Fig. S3G and S3H**). These results suggest that reduced proliferation caused by deactivation of PfNOT1.1 or PfNOT1.2 results from impaired egress and/or invasion process, likely as a reflection of a decreased fitness of the mutant merozoites.

### 4. PfNOT1.1 and PfNOT1.2 have opposing effects on transcript abundance

Given the importance of CCR4-NOT in regulation of gene expression, we explored the role of two PfNOT1 paralogues in maintenance of the global transcriptional pattern across the *P. falciparum* IDC. In particular, we wished to assess the impact of *PfNot1.1* disruption and *PfNot1.2* knockdown on the global mRNA abundance pattern by analyzing the transcriptomes of the parental and transgenic lines. Transcriptional profiles of each parasite line were assessed by microarray at six time points across one IDC in biological triplicates as previously described [4]. Compared to their parental lines, we observed significant changes in mRNA abundance (p < 0.05) with > 1.5 fold-change for 715 genes and 202 genes in ΔPfNot1.1 and ddΔPfNot1.2, respectively (See gene lists in **Table. S3**). A total of 782 genes were significantly affected by either *PfNot1.1* disruption or *PfNot1.2* knockdown across the IDC particularly at the trophozoite and schizont stages (**Fig. 3A**). Interestingly, in ΔPfNot1.1, the majority of mRNA transcripts exhibited decreased levels while in ddΔPfNot1.2 the trend was towards upregulation. The opposing pattern of these transcriptional changes in ΔPfNot1.1 and ddΔPfNot1.2 was also supported by negative Spearman coefficients between the overall transcriptional profiles at each time point (**Fig. 3B**). Indeed, there is a statistically significant overlap (p < 0.001) between these two genes sets with 135 common genes (**Fig. 3C**), 114 of which display opposing changes at the corresponding time points. The changes in mRNA abundance determined by microarray were also validated by quantitative reverse transcriptase-PCR (qRT-PCR) for a random selection of genes at several time points (**Fig. 3D**). Closer inspection of the mean-centered mRNA expression profiles revealed that the apparent transcriptional suppression in ΔPfNot1 (**Fig. 3A, left heat map)** corresponds to delayed profiles of the differentially expressed genes during their active transcription (**Fig. 3A, right panel, green curves).** Conversely, the transcriptional inductions in ddΔPfNot1.2 (**Fig. 3A, right heat map)** correspond to accelerated patterns of the IDC mRNA abundance profiles (**Fig. 3A, right panel, red curves).** Taken together, these observations suggest that PfNOT1.1 and PfNOT1.2 play roles in the timing of the mRNA levels during the “just-in-time”-like transcriptional cascade in the *P. falciparum* IDC during their active transcription (rise of the individual transcript levels prior to their peak abundance). Given that the deletion of PfNOT1.1 caused general down regulations of the transcriptlevels, it is feasible to suggest that the endogenous functions of this paralogues of the CCR4-NOT scaffold subunit contributes to “inductions” of transcript levels; possibly via mRNA stabilization (see below). *Vice versa*, PfNOT1.2 likely contributes to transcriptional “suppressions”.

**Figure 3.**
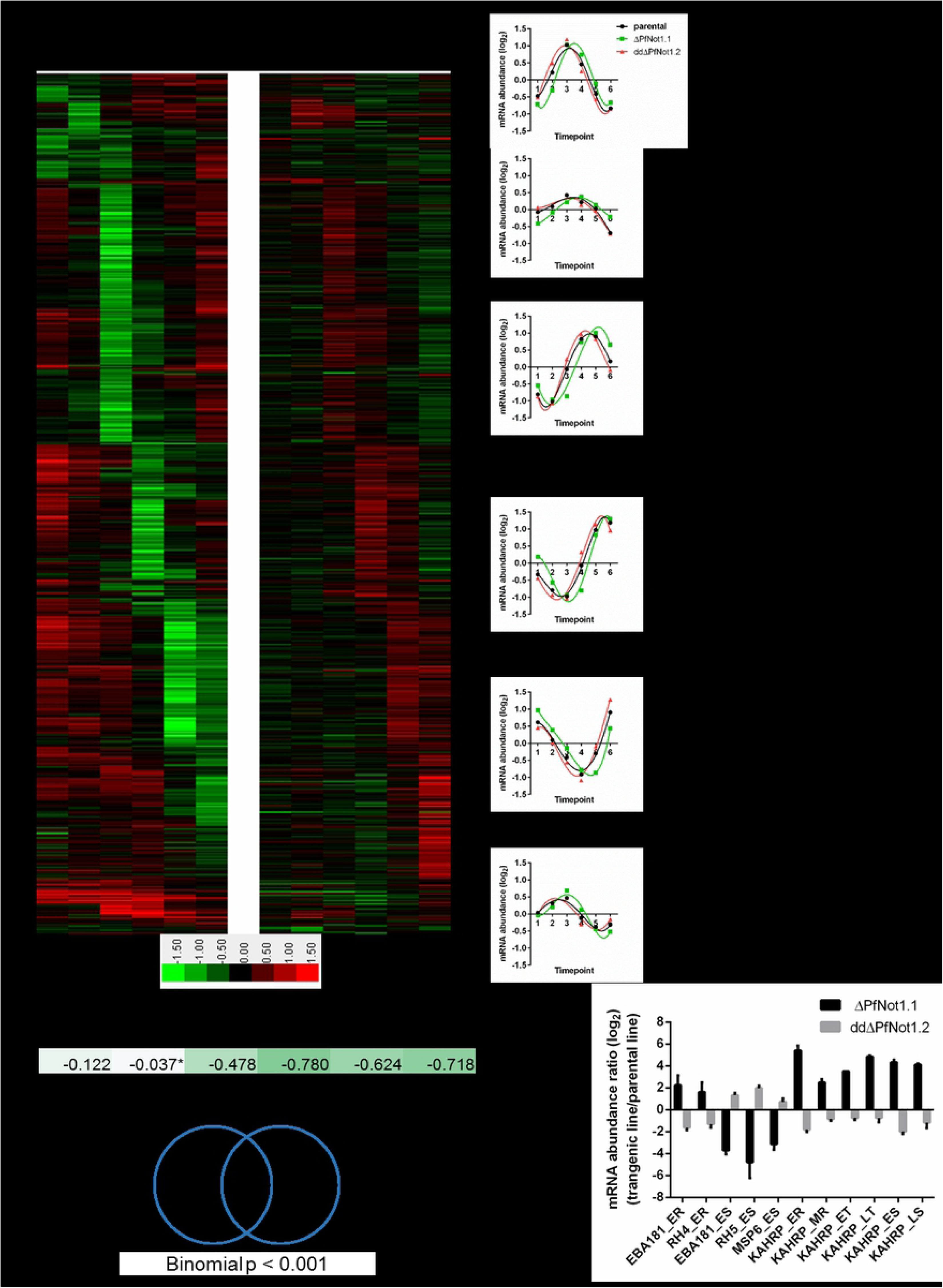
PfNOT1.1 and PfNOT1.2 have opposing effects on transcript abundance. **(A) Left panel.** A total of 782 genes display significantly differential mRNA abundance (> 1.5 fold-change, p < 0.05) in either ΔPfNot1.1 or ddΔPfNot1.2 at six timepoints, representing early ring (ER, 4 hpi), middle ring (MR, 12 hpi), late ring/early trophozoite (ET, 20 hpi), late trophozoite (LT, 28 hpi), early schizont (ES, 36 hpi) and late schizont (LS, 44 hpi). A hierarchical clustering of log_2_-transformed ratios of mRNA abundance in the transgenic lines to that in the parental strain is shown in left panel with 6 clusters categorized based on the timepoint of drastic changes in mRNA abundance. The scale bar at the bottom shows increased abundance in red and decreased abundance in green in the transgenic lines. **Middle panel.** In each cluster, average mean-centered expression of genes at each timepoint was calculated in the parental strain, ΔPfNot1.1 and ddΔPfNot1.2 and then plotted across the IDC. **Right panel.** The enriched GO/MPM pathways in each cluster are listed. **(B)** Spearman coefficient between ΔPfNot1.1 and ddΔPfNot1.2 at each timepoint indicates significant negative correlation at the trophozoite and schizont stages. **(C)** Disruption of *PfNot1.1* or *PfNot1.2* altered mRNA abundance of 715 genes and 202 genes, respectively with a significant overlap of 135 genes (p < 0.001). **(D)** qRT-PCR analysis was performed on random genes at several timepoints in biological triplicates and data are represented as mean ± standard deviation. Changes in transcript abundance were consistent with those assessed by microarray analysis, verifying the accuracy of genome-wide microarray.

Crucially, the PfNOT1-driven transcriptional alterations did not affect the progression of the entire IDC cascade (see **Fig. S3G and S3H**) but was confounded to a distinct subsets of the IDC-driven transcriptionally regulated genes. In order to evaluate the biological relevance of the antagonistic regulatory relationship between PfNOT1.1 and PfNOT1.2 we inspected the functional relevance of each differentially expressed gene cluster (**Fig. 3A right panel**). At the early trophozoite stage, the dysregulated genes are mainly involved in basic cellular and biochemical pathways such as components of apicoplast and mitochondrion, DNA repair, DNA replication, mitosis, transcription, ATP synthesis and metabolic pathways (pyrimidine metabolism, purine metabolism and redox metabolism). Notably, many transcripts encoding egress- and invasion-related genes, merozoite components and merozoite motility proteins were dysregulated at the late trophozoite and the schizont stages, which may lead to the defective invasion rate and subsequent proliferation as observed in ΔPfNot1.1 and ddΔPfNot1.2. Transcriptional changes were also observed in abundant transcripts encoding exported proteins and Maurer-s cleft components, which are associated with erythrocyte remodeling and parasite virulence [54, 55]. Taken together, all the above mentioned affected cellular and biochemical functionalities may represent key processes in the parasite development and thus their dysregulation may lead to suboptimal formation of the invasive merozoites that have a lower potential for successful invasion.

### 5. PfNOT1.1 and PfNOT1.2 regulate gene expression post-transcriptionally

Considering the canonical function of CCR4-NOT complex, the antagonistic activities of PfNOT1.1 and PfNOT1.2 in gene expression may happen at the transcriptional and/or post-transcriptional regulatory steps of mRNA levels. To study this, we explored the effect of the *PfNot1.1* deletion on both transcription and mRNA decay. To study transcription, we assessed the recruitment of the elongating form of RNAPII at six time points across the IDC by chromatin-immunoprecipitation using Ser2/5-P antibody raised previously [10]. Transcriptional status of RNAPII is known to be associated with sequential changes of phosphorylation at the C-terminal domain (CTD) of RPB1, with phosphorylation of Ser5 alone during transcriptional initiation and phosphorylation of both Ser2 and Ser5 (Ser2/5-P) when the polymerase is engaged in elongation [56]. Here, we observed a total of 150 genes that displayed significantly differential enrichment of RPB1 (> 1.5 fold-change, p < 0.05) between ΔPfNot1.1 and its parental strain across the IDC (**Table S4**). In contrast to differential gene expression, the differential RPB1 occupancy in ΔPfNot1.1 was most apparent at ring and trophozoite stages (**Fig. S4A**). Indeed there was a minimal overlap between the 150 genes with differential RPB1 enrichment and the 715 genes with differential mRNA abundance with only 17 genes in both gene sets is statistically insignificant (Binomial p > 0.05) (**Fig. 4A**). Moreover, no Spearman correlation could be observed between RBP1 occupancy and differential mRNA profiles at each time point (**Fig. S4B**). Thus, our data indicates that while deletion of *PfNot1.1* does affect the recruitment of the elongating form of RNAPII, changes in transcriptional elongation are not directly responsible for the altered mRNA abundance of the 715 genes in ΔPfNot1.1.

**Figure 4.**
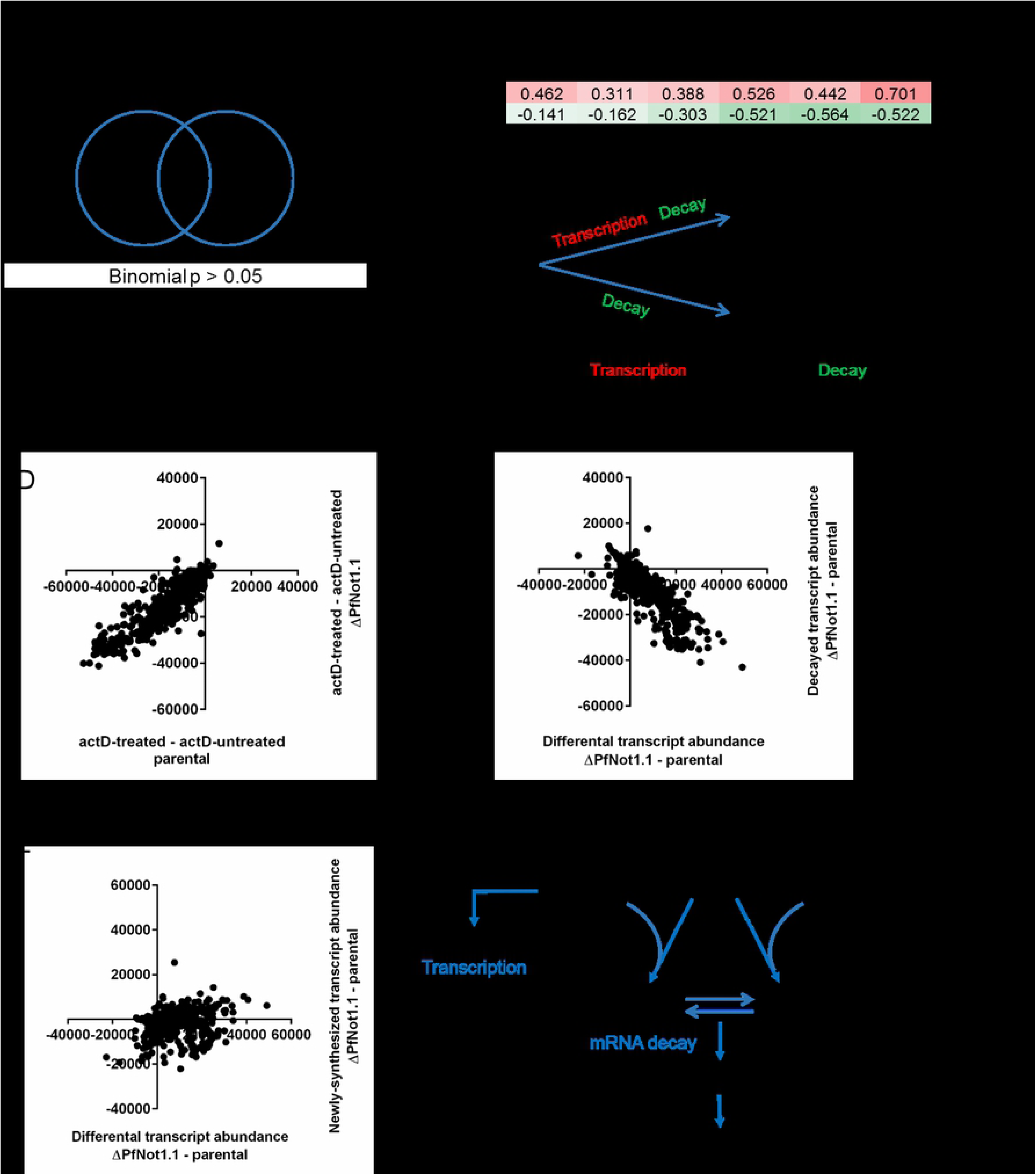
PfNOT1.1 and PfNOT1.2 regulate gene expression post-transcriptionally. **(A)** A total of 150 genes showing significantly differential RNAPII recruitment in ΔPfNot1.1, 17 of which displays significant changes in mRNA abundance (Binomial p > 0.05). **(B)** Spearman correlation coefficient calculated between differential mRNA abundance in three transgenic lines demonstrates a positive correlation between ΔPfNot1.1 and ΔPfCaf1 as well as a negative correlation between ddΔPfNot1.2 and ΔPfCaf1. **(C)** In ΔPfNot1.1 and the parental strain, mRNA decay was assessed using actD or DMSO vehicle followed by microarray analysis. Transcript abundance for each microarray oligo was derived from microarray readings. **(D)** In the presence of actD, transcription inhibition was validated by lower transcript abundance in the actD-treated parasite compared to that in the actD-untreated parasite for both ΔPfNot1.1 and the parental strain. Changes in decayed transcript abundance **(E)** and changes in newly-synthesized transcript abundance **(F)** are plotted against changes in differential transcript abundance between ΔPfNot1.1 and the parental strain within the 4-hour incubation. Changes in transcript abundance are highly correlated to changes in decayed transcript abundance, suggesting that mRNA decay is the key factor that leads to altered transcript abundance in ΔPfNot1.1. **(G)** Proposed mechanism of PfNOT1.1- and PfNOT1.2-mediated gene regulation. PfNOT1.1 and PfNOT1.2 compete to be the scaffold for PfCAF1 to form either a PfNOT1.1-CAF1 complex or PfNOT1.2-CAF1 complex. The two forms function oppositely in regulating mRNA abundance, possibly by controlling PfCAF1 function in mRNA decay. Genes being regulated in this manner are involved in merozoite egress, merozoite invasion, parasite virulence and other cellular processes.

In eukaryotic organisms, NOT1-CAF1 interaction is required for the function of CAF1 for mRNA decay and deadenylation [26–28]. To test whether this association is conserved in *P. falciparum*, we compared significant transcriptional changes of 782 genes observed in ΔPfNot1.1 or ddΔPfNot1.2 to those in a transgenic line with disrupted *PfCaf1* (ΔPfCaf1) reported previously [17]. Strikingly, differential transcript abundance in ΔPfNot1.1 is positively correlated to that of ΔPfCaf1 (**Fig. 4B and Fig. S4C**), demonstrating that functional correlation of PfNOT1 and PfCAF1 is conserved in *P. falciparum*. Conversely, we observed a negative correlation between differential transcript abundance in ddΔPfNot1.2 and ΔPfCaf1, suggesting that the opposite effects of *PfNot1.1* disruption and *PfNot1.2* knockdown on transcript abundance might be due to their opposing roles in modulating PfCAF1 function.

The removal of the mRNA poly(A) tail by CAF1 deadenylase and CCR4 deadenylase is a key step in the initiation of mRNA decay [29]. Therefore, we next assessed the possibility of that PfNOT1 paralogues mediate regulation of mRNA abundance via mRNA decay. Here we utilized the concept of transcription blocking with actD as established previously [57]. Briefly, synchronized cultures of ΔPfNot1.1 and the parental strain were incubated with 20 μg/ml actD or DMSO vehicle for 4 hours at the trophozoite stage (24 hour post-invasion, hpi) in biological duplicates. Subsequently, we carried out comparative analysis of transcriptomes of the resulting parasites, including initial sample at 24 hpi, actD-treated sample and actD-untreated sample **(Fig. 4C).** Inhibition of transcription by actD was validated by lower transcript abundance in the actD-treated parasites compared to that in the actD-untreated parasites for both ΔPfNot1.1 and the parental strain (**Fig. 4D**). We then calculated differential transcript abundance (transcriptional changes between the actD-untreated sample and the initial sample), newly-synthesized transcript abundance (transcriptional change between the actD-untreated sample and the actD-treated sample) and decayed transcript abundance (transcriptional change between the initial sample and the actD-treated sample) within the 4-hour incubation for each parasite line. When comparing the amount of decayed transcript between ΔPfNot1.1 and the parental strain, a lower level of mRNA decay was observed for the majority of transcripts in ΔPfNot1.1 (**Fig. 4E)**, suggesting that the loss of *PfN*ot1.1 does indeed affect mRNA decay. This observation is consistent with the functional correlation of NOT1.1 and CAF1 reported in eukaryotes that NOT1.1-CAF1 interaction is required for the function of PfCAF1 in mRNA decay [26–28].

Furthermore, we observed that the changes in differential transcript abundance between ΔPfNot1.1 and the parental strain are highly correlated with the changes in decayed transcript abundance (**Fig. 4E**, Spearman coefficient = −0.851, p < 0.001), but not correlated with the changes in newly-synthesized transcript abundance (**Fig. 4F**, Spearman coefficient = 0.195, p < 0.05), suggesting that PfNOT1.1 regulates gene expression predominantly by influencing mRNA decay but not transcription at the trophozoite stage.

## Discussion

The existence of one copy of NOT1 as the scaffold of the CCR4-NOT complex has been reported in yeast [58], *Drosophila* [59], human cells [25] and *Trypanosoma brucei*, a parasitic kinetoplastid belonging to the *Euglenozoa* phylum [60]. Two copies of PfNOT1-encoding genes in P. falciparum is contrasting the majority of eukaryotes but is consistent with other *Plasmodium* species, such as *P. yoelii* [18], making it a unique feature in the *Plasmodium* genus. The presence of two copies of NOT1 is the likely result of gene duplication with the insertions of unique amino acid stretches in PfNOT1.2 MIF4G and NOT1 domains evolving with time. NOT1 is an essential gene in yeast [61], human HEK293T cells [62] and Trypanosome [60]. The requirement for two NOT1 paralogues in *Plasmodium* has therefore been unclear. In this study, we successfully produced a full loss-of-function transgenic line for *PfNot1.1* but not *PfNot1.2*. Consistent with this, an attempt to inactivate multiple genes in *P. berghei* was unsuccessful in knocking out *PbNot1.2* [63]. On the other hand, *PfNot1.1* and *PfNot1.2* have been suggested to be dispensable genes in *P. falciparum* given the recovery of *PiggyBac* transposon-insertions into the coding regions of both PfNOT1 paralogues [64]. However, the essential or dispensable character of both genes is presumptive in these two studies given that the failure to inactivate *PbNot1*.2 may be a simple result of a failure of the targeting vector, and *PiggyBac* insertions in the *PfNot1.2* coding region may not have resulted in a complete loss of function. Our data provides direct evidence that PfNOT1.1 is dispensable in parasite viability and that PfNOT1.2 might be refractory to disruption during the IDC (although a possibility of a failure of the targeting vector cannot excluded). These findings raise questions as to why PfNOT1.1 as a more canonical eukaryotic NOT1 is dispensable whereas eukaryotic NOT1 is essential in multiple organisms. Moreover, PfNOT1.2 might possess functions independent from PfNOT1.1 that make PfNOT1.2 refractory to be disrupted. This possibility is supported by the distinct proteins co-immunoprecipitated by PfNOT1.1 and PfNOT1.2 in this study.

The composition of the CCR4-NOT complex is highly conserved across eukaryotes, including NOT1, NOT2, NOT4, NOT5, CAF1, CCR4 and CAF40 [29]. Our Co-IP data suggests that PfCCR4-NOT complexes comprise of either PfNOT1.1 or PfNOT1.2 scaffold in the mutually exclusive manner, PfCAF1, PfCAF1-associating PfCCR4, PfCAF40 and possibly PfNOT4, which associates with PfCAF40 and PfCAF1. In the *P. yoelii* study, PyCCR4 associated with PyNOT1.1, PyNOT1.2, PyCAF1, PyCAF40 and PyNOT2 in schizont parasites. PyCCR4 also co-immunoprecipitated PyNOT5 and PyNOT4 with much lower SAINT stringency [18]. Taken together, the composition of the CCR4-NOT complexes is conserved in *Plasmodium* parasites with NOT4 and NOT5 being less stable components. The weak association of NOT4 with the rest of the complex is expected as NOT4 is a less tightly associated subunit in mammals and *Drosophila* [52, 53]. Contrary to our finding in *P. falciparum*, the association of PyNOT2 with the PyCCR4-NOT complex may result from an additional fixation step the *P. yoelii* study that allows the detection of weakly associated NOT2. Moreover, the weak or missing association of NOT2 and NOT5 with the complex is surprising given the highly conserved interacting domains predicted in both PfNOT1 paralogues, PfNOT2 and PfNOT5. On the other hand, the composition of the CCR4-NOT complex is distinct in another parasite *T. brucei*, in which TbCAF1 co-immunoprecipiated TbNOT1, TbNOT2 and TbNOT5 and no homologues of CCR4 and NOT4 were found in its genome [60]. Notably, the *P. yoelii* study and the *T. brucei* study both reported a punctate distribution of PyCCR4 and TbCAF1 predominantly in the cytoplasm, consistent with our findings on the cytoplasmic localization of four PfCCR4-NOT subunits, suggesting that the post-transcriptional regulatory functions of the CCR4-NOT complex(es) are highly conserved across protozoans though composition varies slightly.

The degradation of mRNA plays important roles in the modulation of gene expression and quality control of mRNA biogenesis. The rate-limiting step of mRNA degradation is deadenylation, the shortening of the poly(A) tail by mRNA deadenylases, including the Pan2/Pan3 complex, the CCR4-NOT complex [23] and the vertebrate-specific DAN/PARN enzyme [65]. CCR4 and CAF1 are the major eukaryotic deadenylases while Pan2/Pan3 and PARN may play secondary roles [66]. Although both CCR4 and CAF1 possess nuclease domain and deadenylase activity [67], studies in yeast suggest that CCR4 is the predominant catalytic subunit of the deadenylase and CAF1 mainly functions to attach CCR4 to the complex [68, 69]. In contrast, CAF1 is suggested to be more important in deadenylation than CCR4 in *C. elegans* [70], *Drosophila* [71] and human HTGM5 fibrosarcoma cells [60]. Similarly, in Trypanosome a protozoan lacking CCR4 homologue, TbCAF1 was shown to function directly in deadenylation in the total RNA and degradation of four mRNAs [60]. In *P. yoelii,* disruption of *PyCcr4* does not affect the length of the poly(A) tail of a PyCCR4-affected/bound transcript in gametocytes, suggesting that CCR4 is not the predominant deadenylase in *Plasmodium* parasites at least in the gametocytes [18]. In the same study, truncated PyCAF1 leads to a comparable but more pronounced effect on parasite transmission compared to the *PyCcr4* disruption, hinting that PyCAF1 plays a more important role in mRNA metabolism as reported in eukaryotes except for yeast. However, the mechanism underlying the function of CAF1 in gene regulation has not been described for either *P. yoelii* [18] or *P. falciparum* [17]. In this study, we showed the functional correlation of PfNOT1.1 and PfCAF1 as well as the effect of disrupted *PfNot1.1* in mRNA decay, providing the first direct evidence on the role of the PfCCR4-NOT complexes in mRNA degradation in *Plasmodium* parasites. In addition, CAF1 plays a structural role linking CCR4 to the rest of the complex and this association is needed for the deadenylation activities of CCR4 [26, 27]. This association is conserved in *Plasmodium* parasites given the similar phenotypes of parasites lacking P*yCaf1* or *PyCcr4* in *P. yeolii* [18]. Our data cannot distinguish whether the effect of *PfNot1.1* disruption on mRNA decay is the direct result of impairment of PfCAF1 activity, or non-exclusively, the indirect effect of altered PfCCR4 attachment to PfCCR4-NOT complexes.

Although NOT1 itself does not contain any nuclease domains, it is associated with deadenylation and mRNA degradation by affecting the function of CAF1 deadenylase through the NOT1-CAF1 interaction [29]. Our data on the physical interaction and the functional correlation of PfNOT1.1 and PfCAF1 indicate that this association in eukaryotes is very likely conserved in the parasite. In the absence of PfNOT1.1, free PfCAF1 deadenylation function is impaired, leading to defective mRNA decay as we observed in ΔPfNot1.1. The decreased mRNA decay affects transcript abundance and results in abnormal timing of gene expression, reflected by the shifted expression pattern we observed in ΔPfNot1.1. On the other hand, the negative functional correlation of PfNOT1.2 and PfCAF1 suggests an inhibitory role of PfNOT1.2 on PfCAF1 function. A reduced protein level of PfNOT1.2 could release PfCAF1 from the PfNOT1.2 complex, allowing it to attach to the PfNOT1.1 complex and function in mRNA degradation, which can be tested by assessing mRNA decay in ddΔPfNot1.2 in future studies. Overall, we propose that PfNOT1.1 and PfNOT1.2 compete to be the scaffold for PfCAF1 binding (**Fig. 4G**). PfNOT1.1 facilitates PfCAF1 function as reported in other eukaryotes, whereas PfNOT1.2 sequesters PfCAF1 and renders it inactive or unavailable. Parasites could favor the formation of one PfNOT1-CAF interaction or the other depending on host response, environmental, metabolic or stage-specific cues, thereby allowing the parasite to modulate gene expression according to circumstance.

In conclusion, we have shown that PfNOT1.1 and PfNOT1.2 are mutually exclusive scaffolds of cytoplasmic CCR4-NOT complexes in *P. falciparum*. PfNOT1.1 and PfNOT1.2 play critical roles in regulating transcript abundance across the IDC. The assessment of the recruitment of the elongating form of RNAPII and mRNA decay in ΔPfNot1.1 underscores the predominant role of PfNOT1.1 in mRNA decay. We propose that PfNOT1.1 and PfNOT1.2 compete to be the scaffold for PfCAF1 binding with PfNOT1.1 facilitating PfCAF1 function and PfNOT1.2 sequestering PfCAF1. Our study significantly extends our knowledge of the CCR4-NOT complex in *Plasmodium* parasites. This study supports the importance of post-transcriptional regulation in *P. falciparum* and provides novel insights into mechanisms of gene regulation in this organism. The unique presence of two PfNOT1 paralogues may open avenues for the development of new drug targets for anti-malarial control.

## Methods

### Malaria Culture

Asexual blood stage of *P. falciparum* T996 strain and transgenic lines were cultured in RPMI-HEPES medium (Gibco), supplemented with 0.25% Albumax (Gibco), 0.2% sodium bicarbonate (Sigma), 100 µM hypoxanthine (Sigma) and 10 µg/ml Gentamycin (Gibco) [72] with 2% hematocrit. Rings were synchronized with 5% sorbitol at 4 hpi and 20 hpi for two continuous IDC to obtain highly synchronized culture [73].

For proliferation rate, parasitemia at the trophozoite stage, starting from 0.1% synchronized culture was measured every IDC using Giemsa-stained thin blood smear. Invasion rate, calculated as the ratio of daughter merozoite parasitemia versus the starting synchronized schizont parasitemia, measures how many merozoites produced from one schizont can successfully invade RBCs. The number of merozoite was counted in 50 late schizonts under microscopy for each parasite line in biological duplicates.

### Creation of transgenic parasite lines

To create transgenic lines with HA-tag in C-terminal endogenous gene, the plasmid was integrated into genome by single crossover recombination. The homology region of PfNOT1.1 and PfNOT4 as a portion of 3’ genomic locus up to but not including the stop codon, was PCR-amplified from T996 genomic DNA and inserted into pTCherryHA plasmid [74] between restriction sites XmaI and BsrGI. The homology region of PfNOT1.2 was inserted into HSP101-HADB plasmid [75] between restriction sites XhoI and EagI to construct pLPfNOT1.2 plasmid. The primers 3HA_F and 3HA_R were annealed by heating at 95°C for 5 min and cooling down at 25°C for one hour. The annealed primer pair was then inserted into pLPfNOT1.2 plasmid between restriction sites XmaI and BglII. The homology region of PfCAF1 was inserted into pTCherryHA plasmid between restriction sites XmaI and NheI.

To disrupt the endogenous *PfNot1.1* by double crossover recombination, two homology regions were inserted into pCC1 plasmid [76] to flank a *hDHFR* cassette between restriction sites SacII and SpeI as well as EcoRI and NcoI to create pCC1-PfNOT1.1KO plasmid. Similarly, pCC1-PfNOT1.2KO plasmid was created to disrupt the endogenous *PfNot1.2*.

To tag the C-terminal endogenous *PfNot1.2* with three HA epitopes and an ecDHFR destabilizing domain, a portion of the 3’ genomic locus of *PfNot1.2* up to but not including the stop codon was amplified and inserted into HSP101-HADB plasmid between restriction sites, XhoI and AvrII, to construct pLPfNOT1.2-HA-ecDDD plasmid. The parasite was co-transfected with a plasmid expressing hDHFR in order to confer resistance to toxicity of the stabilizing ligand, TMP. The parental line was also transfected with the plasmid expressing hDHFR.

*P. falciparum* T996 strain at ring stage was transfected with plasmids by electroporation, established elsewhere [77]. The selection drugs used were 2.5 nM WR99210 (Chemopharma) for the *hDHFR* selective marker, 2.5 μg/ml Blasticidin S (Invitrogen) for the *BSD* selective marker and TMP (Sigma) ligand for the ecDHFR destabilizing domain. Once observing the parasites, the drug cycling was performed to increase the population of parasites with successful integration.

The single clone of parasites was obtained by limiting dilution [78], in which the culture was diluted to a concentration of 0.3 parasite/well in the 96-well plate. The integration of plasmids into the endogenous loci was verified by PCR on genomic DNA extracted using the DNeasy Blood & Tissue Kit (Qiagen).

### Immunoprecipitation and mass spectrometry

Saponin-lysed schizont pellet (400 μl) of the parental line and HA-tagged transgenic lines was lysed with 5 pellet volumes of Lysis Buffer (20 mM HEPES, pH7.8; 10 mM KCl; 1 mM EDTA, pH8.0; 0.65% Nonidet P-40; 1 mM DTT; 1 mM PMSF; 1x protease inhibitor) to obtain cytoplasmic lysate and nucleus pellet, which was then lysed with 2.5 pellet volumes of Extraction Buffer (20 mM HEPES, pH7.8; 400 mM KCl; 1 mM EGTA, pH8.0; 1 mM EDTA, pH8.0; 1 mM DTT; 1 mM PMSF; 1x protease inhibitor) and diluted with 2.5 pellet volumes of Dilution Buffer (20 mM HEPES, pH7.8; 1 mM EDTA, pH8.0; 0.2% Triton X-100; 30% glycerol; 2 mM DTT; 1 mM PMSF; 1x protease inhibitor) to obtain nuclear lysate. Cytoplasmic lysate and nuclear lysate were combined, a portion of which was stored as input lysate and the rest was subjected to immunoprecipitation by HA magnetic bead (Pierce) for 16 hours. Proteins bound on the beads were eluted in 1x Laemmli Buffer supplemented with 0.1 M DTT by boiling at 100°C for 10 min. Prior to mass spectrometry, the efficiency of immunoprecipitation was confirmed by western blotting. The bound proteins in biological duplicates were analyzed by LC-MS/MS mass and the peptides were matched to the PlasmoDB database [79] using Mascot search software with MS tolerance of 10 ppm and MS/MS tolerance of 0.8 Da [80].

### Western blotting

To obtain total protein lysate, saponin-lysed schizont pellet was lysed with 4 pellet volumes of Total Lysis Buffer (4% SDS; 0.5% Triton X-100; 50% 1x PBS) by boiling at 100°C for 10 min. HA-tagged proteins were detected using 1:1,000 anti-HA antibody (Roche) and 1:5,000 anti-rat antibody (Abcam). Actin as a loading control for immunoprecipitation blot was detected using anti-β-actin antibody (Sigma) and 1:2,000 anti-mouse antibody (Cell Signaling). Aldolase as a cytoplasmic marker was detected by 1:2,000 anti-aldolase antibody (Abcam) and 1:2,000 anti-rabbit antibody (GE Health). Histone H3 as a nuclear marker was detected using 1:1,000 anti-histone H3 antibody (Abcam) and 1:2,000 anti-rabbit antibody. The ladder used for proteins larger than 240 kDa was 12 µl HiMark Pre-Stained Protein Standard (Thermo Scientific). The ladder used for proteins smaller than 240 kDa was 2.5 µl BLUltra Prestained Protein Ladder (GeneDireX). The secondary antibodies were conjugated with HRP, signals of which was exposed to an X-ray film (Canon) or captured in ImageQuant LAS4000 Chemiluminescent and Fluorescence Imaging System (Fujitsu Life Sciences) and the quantified using ImageJ [81].

### Immunofluorescence assays

The parental line and HA-tagged transgenic lines were prepared for immunofluorescence assays as previously described [82]. Asynchronous culture in blood smears were fixed in 4% paraformaldehyde and 0.0075% glutaraldehyde for 30 min, permeabilized in 0.1% Triton X-100 for 10 min, blocked in 3% bovine serum albumin for 1 hour, incubated with 1:100 anti-HA antibody and 1:1,000 anti-aldolase antibody overnight at 4°C, and eventually incubated with 1:1,000 DAPI (Thermo scientific), 1:1,000 Alexa Fluor 488 goat anti-rat (Thermo scientific) and 1:1,000 Alexa Fluor 555 goat anti-rabbit (Thermo scientific) in dark for 1 hour. Fluorescence images were taken using a Olympus IX83 Inverted Microsope equipped with a Plan Apo 100x oil objective lens, processed by Metamorph software and merged by ImageJ software.

### Transcriptional profiling by microarray

Total RNA isolated by TRIzol/chloroform method, first strand cDNA synthesis and the subsequent SMART amplification of double-stranded cDNA were established elsewhere [83]. For each microarray hybridization, 4 µg sample cDNA was coupled with Cy5-dye and 4 µg reference cDNA, a pooling of 3D7 parasite harvested at 8 timepoints across IDC, was coupled with Cy3-dye. The microarray chip, containing 11,005 probes covering 5,061 genes in *P. falciparum* genome, was scanned by Tecan PowerScanner and oligos with bad signals were flagged for deletion. Background subtraction, normalization and filtering of raw signal intensities were accomplished by the “limma” package in R [84]. Expression of each gene was averaged from the log_2_-transformed expression of oligos for the specific gene. The parasite age for each sample was subsequently estimated with respect to a reference IDC transcriptome (T996 strain) based on a Spearman Rank Correlation Coefficients as previously estabilished [85] and modified [86].

Genes showing significantly differential transcript abundance between the transgenic lines and the parental strain were filtered with a fold change cutoff greater than 1.5, p < 0.05 (pairwise Student’s t-test). Hierarchical clustering was carried out using Cluster 3.0 [87] and visualized using Java Treeview [88]. Raw microarray data was submitted to the NCBI GEO public database. Gene set enrichment analysis [89] was carried out on genes with differential transcript abundance to find out enriched functional pathways. Correlation of differential transcript abundance in different parasite lines was assessed based on Spearman Rank Correlation Coefficient.

### Quantitative real time PCR

Extracted total RNA was reverse transcribed into cDNA using SuperScript III first-strand synthesis system for RT-PCR (Invitrogen). Transcript levels of selected targets and the internal control, serine-tRNA ligase, in 20 ng cDNA were measured by real-time PCR in triplicates using SYBR Select Master Mix (Thermo Scientific). The log_2_-transformed fold-change of transcript abundance (-ΔΔC_T_) was calculated using the equation:

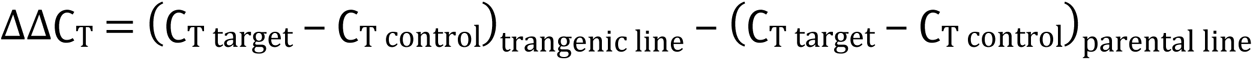

### Chromatin-immunoprecipitation

At 6 timepoints across one IDC, 1.5 ml iRBCs with 5% parasitemia were crosslinked with 0.1% formaldehyde and chromatin immunoprecipitation was performed as previously described [83]. Briefly, parasite was lysed with Buffer A (1 mM EDTA, pH8.0; 0.25% NP40; 1x protease inhibitor; 2 mM PMSF) and the resulting nuclei pellet was lysed with SDS Lysis Buffer (1% SDS; 50 mM Tris, pH8.1; 10 mM EDTA; 1x protease inhibitor; 2 mM PMSF), sonicated into 200-1,000 bp DNA fragments and diluted with ChIP Dilution Buffer (0.01% SDS; 1.1% Triton X-100; 16.7 mM Tris, pH8.1; 1.2 mM EDTA; 167 mM NaCl; 1x protease inhibitor; 2 mM PMSF). A portion of chromatin DNA was stored for input DNA and the rest was incubated with protein A agarose /salmon DNA beads for 2 hours aiming to reduce unspecific binding and then incubated with anti-Ser2/5-P antibody for 16 hours. The antibody:protein was immunoprecipitated by pre-blocked protein A agarose/salmon DNA beads and eluted. The beads were washed twice with Low Salt Wash Buffer (0.1% SDS; 2 mM EDTA; 20 mM Tris-Cl, Ph8.1; 150 mM NaCl), thrice times with High Salt Wash Buffer (0.1% SDS; 2 mM EDTA; 20 mM Tris-Cl, Ph8.1; 500 mM NaCl), once with LiCl Wash Buffer (0.25 M LiCl; 1% NP-40; 1% deoxycholic acid; 1 mM EDTA; 10 mM Tris-Cl, pH8.1) and twice with TE Buffer (10 mM Tris-Cl, pH8.1; 1 mM EDTA). The antibody:protein was eluted in Elution Buffer (1% SDS; 0.1 M sodium bicarbonate). ChIP DNA and input DNA were reverse-crosslinked, amplified and hybridized on microarray chip in biological triplicates. A fold-change cutoff more than 1.5, p < 0.05 (pairwise Student’s t-test) was employed to filter genes with significantly differential RPB1 recruitment.

### mRNA decay assay

To stop transcription, the synchronized ΔPfNot1.1 and the parental strain at 24 hpi was incubated in 20 μg/ml actD (Thermo Scientific) or DMSO for 4 hours, as established previously [57]. For each parasite line, the initial sample at 24 hpi, the actD-treated sample at 28 hpi and the actD-untreated sample at 28 hpi were harvested in biological replicates. Transcriptional profile of each sample was assessed by microarray as described above. Transcript abundance is represented as the cy5-channel reading of 11,005 microarray oligo, which was normalized to the microarray scanning intensity. For each parasite line within the 4-hour incubation, the amount of differential transcript abundance, decayed transcript and newly-synthesized transcript were calculated. Changes in these parameters were compared between ΔPfNot1.1 and the parental strain.

### Primer list

The primers are in the 5’- to 3’- orientation. The 21-bp extension for ligation-independent cloning, described elsewhere [90] is indicated in the small case.

pTPfNOT1.1-3HA_HR_F: aacaaatatagatctcccgggTTCATAACTCTTCACAATCACATC

pTPfNOT1.1-3HA_HR_R: gtacgggtaaagcttgtacagTTCTTTAAATGTTAAATTTTTTTTTTTCTGG

pTPfNOT4-3HA_HR_F: aacaaatatagatctcccgggAGATGATGAAGATGAAGAAGATGAC

pTPfNOT4-3HA_HR_R: gtacgggtaaagcttgtacagTGCACTTAAATTTAGCTTCTCATTC

pLPfNOT1.2-3HA_HR_F: gacactatagaactcgagTTCATGGGTTGAACTTATATCATC

pLPfNOT1.2-3HA_HR_R: ttatataactcgacgcggccgagatctcagtcccgggAACGCGCCCTAGGCATGCTTG

3HA_F:

CCGGGTACCCATACGATGTTCCAGATTACGCTTACCCATACGATGTTCCAGATTACGCTTACCCATACGATGTTCCAGATTACGC TTGAA

3HA_R:

GATCTTCAAGCGTAATCTGGAACATCGTATGGGTAAGCGTAATCTGGAACATCGTATGGGTAAGCGTAATCTGGAACATCGTA TGGGTAC

pTPfCAF1-3HA_HR_F: atctcccgggAATGAAGGAATTATTATGCATG

pTPfCAF1-3HA_HR_R: gccatgctagcGTTATCATAAAAATATTTATGGTCCTTAG

pCC1-PfNOT1.1KO_HR1_F: tccaatggcccctttccgcggGATCGAAAAATAATGATGTG

pCC1-PfNOT1.1KO_HR1_R: aagacagatcttcgtactagtTTTGGTACTTGTTGTTGATA

pCC1-PfNOT1.1KO_HR2_F: tatttattaaatctagaattcCTCATCCATAATTATAGGAGAAG

pCC1-PfNOT1.1KO_HR2_R: gccagcctaggagttccatggAGTTCCTCTAATTTCGTACTATC

pCC1-PfNOT1.2KO_HR1_F: tccaatggcccctttccgcggGAGGATGGATTACCATGA

pCC1-PfNOT1.2KO_HR1_R: aagacagatcttcgtactagtGAATGTGTTCATTTTCACTT

pCC1-PfNOT1.2KO_HR2_F: tatttattaaatctagaattcGATATGGTAGATACAAATGGAG

pCC1-PfNOT1.2KO_HR2_R: gccagcctaggagttccatggCTTATGTGATCTTGATTCTTC

pLPfNOT1.2-3HA-ecDDD_HR_F: gacactatagaactcgagTTCATGGGTTGAACTTATATCATC

pLPfNOT1.2-3HA-ecDDD_HR_R: aacatcgtatgggtacctaggAACGCGCCCTAGGCATGCTTG

PfNOT1.1-HA_1: ATATATAGGAATGTCATTACCATC

PfNOT1.1-HA _2: GTGTATATATAAACACAAGATGTG

PfNOT1.1-HA _3: TAATATAGAAAAAGTTACATGCACG

PfNOT1.1-HA _4: CCTAAGTCATTAAGGTACTAACTTAG

PfNOT1.2-HA _1: CTTTTATTATCACCTTTAAGAATTCCTG

PfNOT1.2-HA _2: GTTAAATATTTAATTAGGAAAGGATAACATG

PfNOT1.2-HA _3: GTATATTGGGGTGATGATAAAATGAAAG

PfNOT1.2-HA _4: AGTGCCACCTGACGTCTAAGAAAC

PfCAF1-HA _1: GAGGACCAACAAGAAGTAGATC

PfCAF1-HA _2: CATATACATTATTATGTGTGAGTTC

PfCAF1-HA _3: TAATATAGAAAAAGTTACATGCACG

PfCAF1-HA _4: CCTAAGTCATTAAGGTACTAACTTAG

PfNOT4-HA _1: ATGATAATGAAGAAGATCAAGAG

PfNOT4-HA _2: ATATGTCACTTTGTTTTTGTTTC

PfNOT4-HA _3: TAATATAGAAAAAGTTACATGCACG

PfNOT4-HA _4: CCTAAGTCATTAAGGTACTAACTTAG

PfNOT1.1KO_5: ATGTACAAATGGATGAAGAATCTG

PfNOT1.1KO_6: ATATTCTTGACATGTTCCTTTATGC

PfNOT1.1KO_7: TTACCAAATAATCTGAACGCAAAC

PfNOT1.1KO_8: AGATTCTATAGTATTTAAGGATGGTTG

PfNOT1.1KO_9: TTTTTTCTTTTATCATGCACATTGG

PfNOT1.1KO_10: GAAGTATATGAGAAGAATGATTAAGC

PfNOT1.1KO_11: TATATCCAATGGCCCCTTTC

PfNOT1.1KO_12: GCGGTGTGAAATACCGCAC

PfNOT1.2KO_5: GAAGAAAAGAAAAAGCAAGGAG

PfNOT1.2KO_6: TTTTATTCGTCACACTATCATTTG

PfNOT1.2KO_7: ATATGAACAGTATCAACAATATGAAC

PfNOT1.2KO_8: TGATTCATATGTTTTTCATCTTCTAC

PfNOT1.2KO_9: CAAAATGCTTAAGACAGATCTTC

PfNOT1.2KO_10: CTTTTACAATATGAACATAAAGTACAAC

PfNOT1.2KO_11: TATATCCAATGGCCCCTTTC

PfNOT1.2KO_12: GCGGTGTGAAATACCGCAC

PfNOT1.2-HA-ecDDD_13: CTTTTATTATCACCTTTAAGAATTCCTG

PfNOT1.2-HA-ecDDD_14: GTTAAATATTTAATTAGGAAAGGATAACATG

PfNOT1.2-HA-ecDDD_15: TTTCCATGCCGATAACGTGATCTAC

PfNOT1.2-HA-ecDDD_16: AGTGCCACCTGACGTCTAAGAAAC

The reverse primer contains the promoter sequence of T7 RNA polymerase for in-vitro transcription, shown in small cases.

PfNOT1.1_probe 1_F: ACCAGTAGCTGTCGATAGAGC

PfNOT1.1_probe 1_R: actgactgtaatacgactcactatagggACGTAATGGTTCTTTGCATGTAGC

PfNOT1.1_probe 2_F: CATCAGAAACAAATATCATCGATCC

PfNOT1.1_probe 2_F: actgactgtaatacgactcactataggggGTGTATATATAAACACAAGATGTG

PfNOT1.1_probe control_F: TGGAAGAATGTGCCGAACTTG

PfNOT1.1_probe control_R: actgactgtaatacgactcactatagggGCACATAATGGTTGTTCTGAGGT

### Data availability

Microarray data is available in GEO database: transcriptional profiling of ΔPfNot1.1 and ddΔPfNot1.2 (GSE142275); recruitment of RNAPII in ΔPfNot1.1 (GSE142276); mRNA decay analysis in ΔPfNot1.1 (GSE142527). All data has been deposited in two datasets in Mendeley Data Repository (DOI: 10.17632/gnytrrsjcn.1).

## Acknowledgment

Y. L. is a recipient of Nanyang Graduate Scholarship from Nanyang Technological University, Singapore. This work was supported by an AcRF Tier 1 grant from the Singapore Ministry of Education (#RG41/09) and grants from the Singapore National Medical Research Council (#NMRC/CBRG/0029/2013; CBRG12nov104) to MF; and AcRF Tier 2 grant from the Singapore Ministry of Education (#MOE2017-T2-2-030 (S)) and Singapore National Medical Research Council (#NMRC/OFIRG/0040/2017) to ZB. We would like to thank Brendan Crabb (pTCherryHA), Alan Cowman (pCC1) and Daniel Goldberg (HSP101-HADB) labs for sharing plasmid vectors. We also thank Siu Kwan Sze lab for performing mass spectrometry.

## Author Contribution

Conceptualization: Y.L., Z.B. and M.F.; Formal Analysis: Y.L.; Funding Acquisition: Z.B. and M.F.; Investigation: Y.L.; Methodology: Y.L. and R.R.; Project Administration: C.Q.Z. and F.R.; Software: L.Z.; Supervision: Z.B. and M.F.; Writing – Original Draft Preparation: Y.L.; Writing – Review & Editing: Y.L., Z.B. and M.F..

The funders had no role in study design, data collection and analysis, decision to publish, or preparation of the manuscript.

## Supporting information captions

**Figure S1. Conserved domains of the NOT1 orthologues in different organisms. (A)** The length and location of four conserved domains, including DUF2363, MIF4G, DUF3819 and NOT1, are proportional to the length of protein sequence. **(B)** The conserved domains were pair-wise blasted using the NCBI BLASTp program. The lengths and the similarities of the identified regions were calculated. Due to the insertions in the MIF4G domain and the NOT1 domain of PfNOT1.2, two parts were blasted separately.

**Figure S2. The CCR4-NOT subunits were successfully tagged with HA epitopes. (A)** The endogenous gene was fused with three HA epitopes by single crossover recombination. **(B-E)** The integration of plasmids was verified by genotyping PCR. Parental loci (par) were detected in the parental lines but not in the transgenic lines, confirming the cloned parasites does not contain contamination of the parental lines. Integrated loci (int) and episomal loci (epi) were detected in the transgenic lines. **(F)** The parental strain was stained with anti-HA antibody, serving as a negative control of the immunofluorescent assay. Nuclei were stained with DAPI and aldolase immunoreactivity was used as a cytoplasmic maker. No signals were detected by anti-HA antibody, confirming that the detected signals in **Fig. 1E-H** are specific. Scale bars are 1 micron.

**Figure S3. Genotyping PCR and additional phenotypes of ΔPfNot1.1 and ddΔPfNot1.2.** Genotyping PCR was performed to verify the integration of the plasmids into the genome in the **(A)** *PfNot1.1* knockout transgenic line, **(B)** *PfNot1.2* knockout transgenic line and **(C)** *PfNot1.2* conditional knockdown transgenic line. Detected genomic loci include parental locus (par, par1 and par2 flanking two homology regions, respectively), integrated locus (int, int1 and int2) and episomal locus (epi, epi1 and epi2). **(D)** Representative western blotting of HA-ecDDD-tagged PfNOT1.2 displayed reduction of protein expression level after removing TMP for three cycles. The failure to detect the full-length NOT1.2-HA-ecDDD protein at 542 kDa is possibly due to the incomplete stabilization of the fusion protein by TMP. The sizes of the ladder are given at the left. **(E, F)** The number of merozoite in each schizont, counted in 50 iRBCs in biological duplicates, is comparable between the transgenic lines and the parental strain. Data are represented as mean ± standard deviation. Statistic test was performed at each timepoint using pairwise Student’s t-test with p > 0.05 shown as non-significant (ns). **(G, H)** Estimated developmental ages of parasites harvested at 6 timepoints across the IDC were comparable between the transgenic lines and the parental strain at biological triplicates. Data are represented as mean ± standard deviation. Statistic test was performed at each timepoint using pairwise Student’s t-test with p < 0.05 (*), which was not applicable for identical ages for triplicates (na).

**Figure S4. PfNOT1.1 regulates transcript abundance by affecting post-transcriptional regulation instead of transcriptional regulation. (A)** Heatmap of differential RNAPII recruitment for 150 genes (left panel) and corresponding differential mRNA abundance (right panel) across the IDC are represented by log_2_-transformed differences in ΔPfNot1.1 versus the parental strain. The scale bar at the bottom shows increased abundance in red and decreased abundance in green in the transgenic lines. **(B)** The Spearman correlation coefficient was calculated between differential RNAPII recruitment and differential mRNA abundance of 150 genes in ΔPfNot1.1 at each timepoint, showing no correlations. **(C)** Significantly differential transcript abundance of 782 genes in ΔPfNot1.1 or ddΔPfNot1.2 is shown as the log_2_-transformed ratio of expression in the transgenic lines versus the parental strain. Differential mRNA abundance of the corresponding genes in ΔPfCaf1 was adapted [17]. The scale-bar is given at the bottom.

**Table S2. List of proteins co-immunoprecipitated by HA-tagged PfNOT1.1, PfNOT1.2, PfCAF1 and PfNOT4.** HA-tagged proteins were co-immunoprecipitated by Pierce HA magnetic beads and bound proteins were identified by LC-MS/MS mass spectrometry in the transgenic lines and the parental strain at the schizont stage in biological replicates. The mass spectrometry data was matched to the PlasmoDB database using Mascot search software and the sequenced peptides were identified based on Mascot probability scoring [80]. The ions score was calculated for each peptide to measure how well the peptide sequence matches the database sequences. The Mascot protein score, as the summed ions score of all peptides identified in one protein reflects the confidence of this protein detected in the eluent. The proteins detected only in the transgenic lines or detected in both the parental strain and the transgenic lines with a log_2_-transformed ratio of Mascot score in the transgenic lines versus that in the parental strain more than 1.5, were selected as positive hits. The proteins identified as positive hits in both biological replicates are listed in this file and ranked based on the average Mascot score.

**Table S3. List of genes with significantly differential mRNA abundance in ΔPfNot1.1 and ddΔPfNot1.2.** The differential mRNA abundance is represented by the log_2_-transformed ratio of mRNA abundance in the transgenic lines versus that in the parental strain. The significant change is defined as an expression fold-change more than 1.5 (corresponding to the log_2_ ratio at 0.585) and p-value (pairwise Student’s t-test) less than 0.05. In ΔPfNot1.1 and ddΔPfNot1.2, there are 715 genes and 202 genes significantly differentially expressed, respectively. Six clusters of genes, categorized according to the timepoint of the most drastic changes in mRNA abundance are also indicated. A total of 782 genes with significant changes in mRNA abundance in either of two transgenic lines and a total of 135 genes showing significantly differential mRNA abundance in both ΔPfNot1.1 and ddΔPfNot1.2 are listed as well.

**Table S4. List of genes with significantly differential recruitment of RNAPII in ΔPfNot1.1**. Recruitment of the RPB1 with phosphorylated Ser2 and Ser5 was assessed by chromatin-immunoprecipitation followed by microarray analysis. The differential recruitment was calculated between ΔPfNot1.1 and the parental strain at six timepoints across the IDC. The significant change is defined as a fold-change more than 1.5 and p-value (pairwise Student’s t-test) less than 0.05. The values are given in the form of log_2_-transformed differential recruitment. The order of the genes corresponds to the heatmap in Fig. S4A in supplemental figures.

